# QTL mapping reveals complex genetic architecture of quantitative virulence in the wheat pathogen *Zymoseptoria tritici*

**DOI:** 10.1101/051169

**Authors:** E. L. Stewart, D. Croll, M. H. Lendenmann, A. Sanchez Vallet, F. E. Hartmann, J Palma Guerrero, Z. Ma, B. A. McDonald

## Abstract

We conducted a comprehensive analysis of virulence in the fungal wheat pathogen *Zymoseptoria tritici* using QTL mapping. High throughput phenotyping based on automated image analysis allowed measurement of pathogen virulence on a scale and with a precision that was not previously possible. Across two mapping populations encompassing more than 520 progeny, 540,710 pycnidia were counted and their sizes and grey values were measured, yielding over 1.6 million phenotypes associated with pathogen reproduction. Large pycnidia were shown to produce more numerous and larger spores than small pycnidia. Precise measures of percent leaf area covered by lesions provided a quantitative measure of host damage. Combining these large and accurate phenotype datasets with a dense panel of RADseq genetic markers enabled us to genetically dissect pathogen virulence into components related to host damage and components related to pathogen reproduction. We show that different components of virulence can be under separate genetic control. Large-and small-effect QTLs were identified for all traits, with some QTLs specific to mapping populations, cultivars and traits and other QTLs shared among traits within the same mapping population. We associated the presence or absence of accessory chromosomes with several virulence traits, providing the first evidence for an important function associated with accessory chromosomes in this organism. A large-effect QTL involved in host specialization was identified on chromosome 7, leading to identification of candidate genes having a large effect on virulence.

## Introduction

Much plant pathology research has been oriented around understanding the classic gene-for-gene (GFG) interaction (Flor 1955) between the pathogen and its host, largely because early plant breeding programs focused on the introgression of major, qualitative resistance genes (R-genes) to control plant pathogens. Because most pathogens evolved quickly to defeat major R-genes, and quantitative resistance appeared more durable, the genetic basis of quantitative resistance became an area of intensive investigation, leading eventually to identification of host genes encoding quantitative resistance (Krattinger et al. 2009; St.Clair 2010). The term aggressiveness is typically used by plant pathologists to describe the quantitative degree of damage caused by a pathogen strain. In the broader literature, virulence is defined as the quantitative degree of damage caused by a pathogen to its host. We will use the latter definition to be consistent with the wider field of life sciences. In contrast to quantitative resistance in the host, quantitative virulence in pathogens is much less studied and remains poorly understood (Lannou 2011; Pariaud et al. 2009), though quantitative virulence appears to be common in plant pathogens (Pariaud et al. 2009).

Given the broad spectrum in observed virulence phenotypes, the genetic architecture of virulence may be mediated both by single genes of large effect that follow the GFG model (Flor 1955) and by many genes that make smaller contributions to quantitative virulence. Quantitative traits often exhibit a continuous distribution in a population and are typically governed by independent assortment of multiple alleles with small individual effects, with different combinations of alleles responsible for the range of observed phenotypes in recombining populations (Mackay 2001). Numerous confounding factors such as dominance, pleiotropy and additive allele effects also contribute to the genetic architecture of a trait (Hansen 2006). In order to disentangle individual loci contributing to overall virulence phenotypes, we utilized quantitative trait locus (QTL) mapping, an established technique for determining the genetic architecture of quantitative traits. QTL mapping was successfully used to identify candidate genes underlying traits of interest in plants (Alonso-Blanco et al. 2014), animals (Solberg Woods 2014) and humans (Almasy et al. 2009). It has also been employed in fungi (Larraya et al. 2003), though its use in fungi remains rare compared to other eukaryotes (Foulongne-Oriol 2012).

Virulence in the wheat pathogen *Zymoseptoria tritici* (formerly *Mycosphaerella graminicola*), causal agent of Septoria tritici blotch (STB), is a predominantly quantitative trait (Stewart et al. 2014; Zhan et al. 2007). *Z. tritici* is the most damaging wheat pathogen in Europe (Jørgensen et al. 2014; O Driscoll et al. 2014) and is considered an important fungal pathogen worldwide (Dean et al. 2012). Under optimal conditions, yield losses can reach 50% (Eyal et al. 1987). Even with the use of resistant cultivars and regular fungicide treatments, yield losses of 5-10% can be expected (Fones et al. 2015). Ten to fourteen days after infection chlorotic areas begin to appear that later become necrotic lesions. Pycnidia containing asexual pycnidiospores develop mostly within these necrotic lesions in the sub-stomatal cavities. Pycnidiospores are exuded from the pycnidia in a gelatinous cirrhus during periods of high humidity and are spread throughout the plant canopy by rain splash. Numerous asexual infection cycles occurring during a growing season are the main cause of STB epidemics in the field.

Despite its agricultural importance, the mechanisms responsible for virulence in *Z. tritici* and their underlying genetics remain poorly understood. Nineteen genes affecting virulence in *Z. tritici* have been described and functionally characterized (Table S1). Most of these genes have developmental regulatory functions or affect inherent fitness without having a specific role in virulence. Host resistance is also quantitative, with 21 genes and 89 genomic regions implicated in resistance to *Z. tritici* (Brown et al. 2015). The quantitative nature of virulence in *Z. tritici* makes QTL mapping an ideal technique to elucidate the genetic determinants of quantitative virulence, as already validated using several other quantitative traits in *Z. tritici* (Lendenmann et al. 2014; Lendenmann et al. 2015; Lendenmann et al. 2016).

Epidemic development is strongly affected by the ability of a pathogen to reproduce and cause subsequent infections (Parlevliet 1979), hence the reproductive output of a pathogen is an important determinant of the total damage it can cause during an epidemic. Damage caused by plant pathogens is frequently assessed by quantifying the area covered by disease lesions on the host. However, a pathogen’s ability to induce lesions does not necessarily reflect its ability to reproduce. Indeed, several studies have shown that pathogen damage, as indicated by the plant area covered by disease lesions, can be independent of pathogen reproduction, as indicated by the number of spores produced during an infection (Bruns et al. 2014; Habgood 1977; Halama et al. 1999; Pariaud et al. 2009; Twizeyimana et al. 2014), suggesting that these traits may be under separate genetic control. The contribution of each virulence component combines to produce quantitative disease phenotypes.

The success of a QTL mapping project depends on the availability of genetic markers and accurate phenotypic data. With current availability of genomic tools that can generate very large numbers of genetic markers comparatively easily and cheaply, the limiting factor in most studies is accurate phenotyping (Furbank et al. 2011). Using image analysis to phenotype *Z. tritici* infection allows for measurement of both leaf damage, based on the percentage of the leaf covered by necrotic lesions, and pathogen reproduction, based on the number and size of the pycnidia formed on the infected leaves (Stewart and McDonald 2014, Stewart et al, 2016).

The aim of this study was to investigate the genetic architecture of virulence traits in *Z. tritici*. We conducted a QTL mapping study using two mapping populations derived from parental isolates exhibiting varying degrees of virulence. Combining digital image analysis with a large number of RADseq SNP markers in a large number of offspring allowed us to genetically separate traits related to host damage and pathogen reproduction. We also identified novel candidate genes involved in host specialization and virulence.

## Results

### Phenotyping

A phenotyping method based on digital image analysis (Stewart et al. 2014) was used to measure % leaf area covered by lesions (PLACL), pycnidia density (# of pycnidia per cm^2^ leaf), pycnidia size and pycnidia melanization in two *Z. tritici* mapping populations (3D1×3D7 and 1A5×1E4) that have also been used to study melanization (Lendenmann et al. 2014), fungicide sensitivity (Lendenmann et al. 2015) and temperature sensitivity (Lendenmann et al. 2016). The pairs of parental isolates used to make the crosses were chosen based on differing levels of virulence observed in previous work (Zhan et al. 2005). To investigate the effect of host on virulence, we used cultivars Runal (moderately susceptible to *Z. tritici*) and Titlis (moderately resistant) (Courvoisier et al. 2015). All phenotypes showed a continuous distribution in both crosses. Transgressive segregation was evident for all phenotypes, with some progeny exhibiting more extreme phenotypes than the parent isolates (Figure 1). In cross 3D1×3D7 all phenotypes were significantly higher on Runal than Titlis, including mean pycnidia size (F=3.93, p=0.048), pycnidia melanization (F=22.7, p<0.001), PLACL (F=21.4, p<0.001), and pycnidia density (F=19.2, p<0.001). In cross 1A5×1E4, PLACL (F=22.7, p<0.001) and pycnidia melanization (F=17.7, p<0.001) were significantly higher in Runal whereas pycnidia size (F=11.5, p<0.001) and pycnidia density (F=73.6, p<0.001) were significantly higher in Titlis. Across both crosses, a total of 540,710 pycnidia were counted and their sizes and melanization measured, yielding over 1.6 million phenotypic data points associated with fungal fruiting bodies. In order to evaluate whether counting and measuring pycnidia provides an accurate estimate of spore number, the spore output of a subset of isolates was measured. The mean number of spores per pycnidium was 2598. The mean spore size, as measured by length, was 17.6 *μ*m. There were weak but significant positive correlations between pycnidia size and spore size (r2=0.111, p=0.001) and between pycnidia size and the number of spores per pycnidium (r2= 0.112, p=0.001). These findings indicate that pycnidia size is correlated with reproductive output of *Z. tritici*.

**Figure 1.1:**
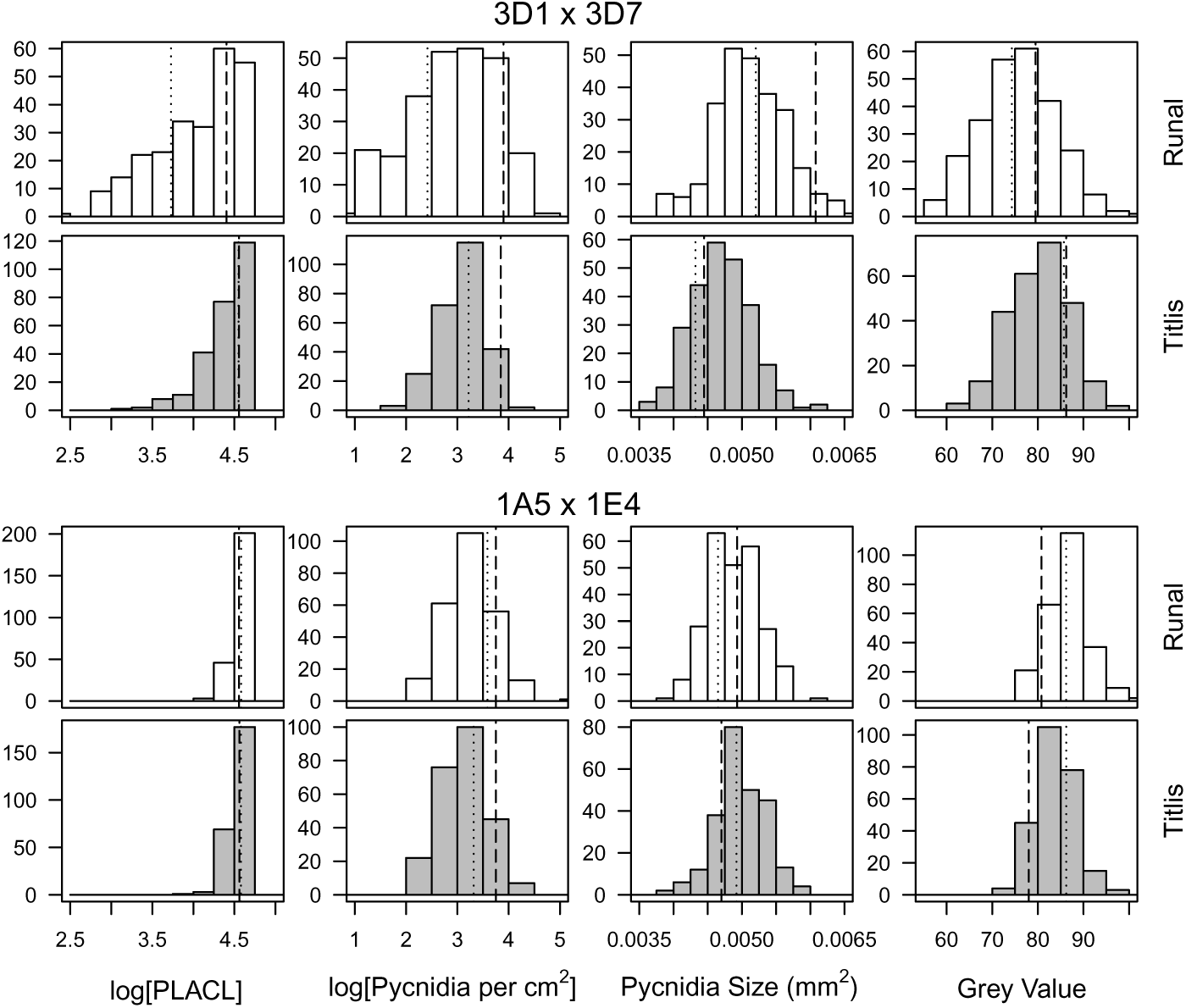
Frequency histograms of virulence phenotypes from the *Zymoseptoria tritici* mapping populations 3D1×3D7 (upper panels) and 1A5×1E4 (lower panels) on the wheat cultivars Runal (white bars) and Titlis (grey bars). Vertical lines represent phenotype values of the parent isolates, dotted line represents 3D1 and 1A5, dashed line represents 3D7 and 1E4.

### Quantitative Trait Locus (QTL) mapping

We used QTL mapping to elucidate the genetic architecture of the quantitative virulence phenotypes. QTLs were identified for all phenotypes. All QTLs in cross 3D1×3D7 were found only on Runal and mapped to a single large effect QTL on chromosome 7 (Figure 2a). In 1A5×1E4 the same QTL for pycnidia density was found on chromosome 5 on both cultivars. A second QTL for pycnidia density was found on chromosome 9 in Titlis and on chromosome 3 in Runal. PLACL in Titlis and pycnidia grey value in Runal both mapped to the same QTL on chromosome 5. Pycnidia size also mapped to the same QTL on chromosome 3 in cv Runal (Figure 2b). No QTLs were shared between the two crosses. The 95% confidence interval for a QTL ranged from 91.3 kb for the chromosome 5 QTL in 1A5×1E4 to 717.7 kb for the chromosome 7 QTL for grey value on Runal in 3D1×3D7.

**Figure 1.2:**
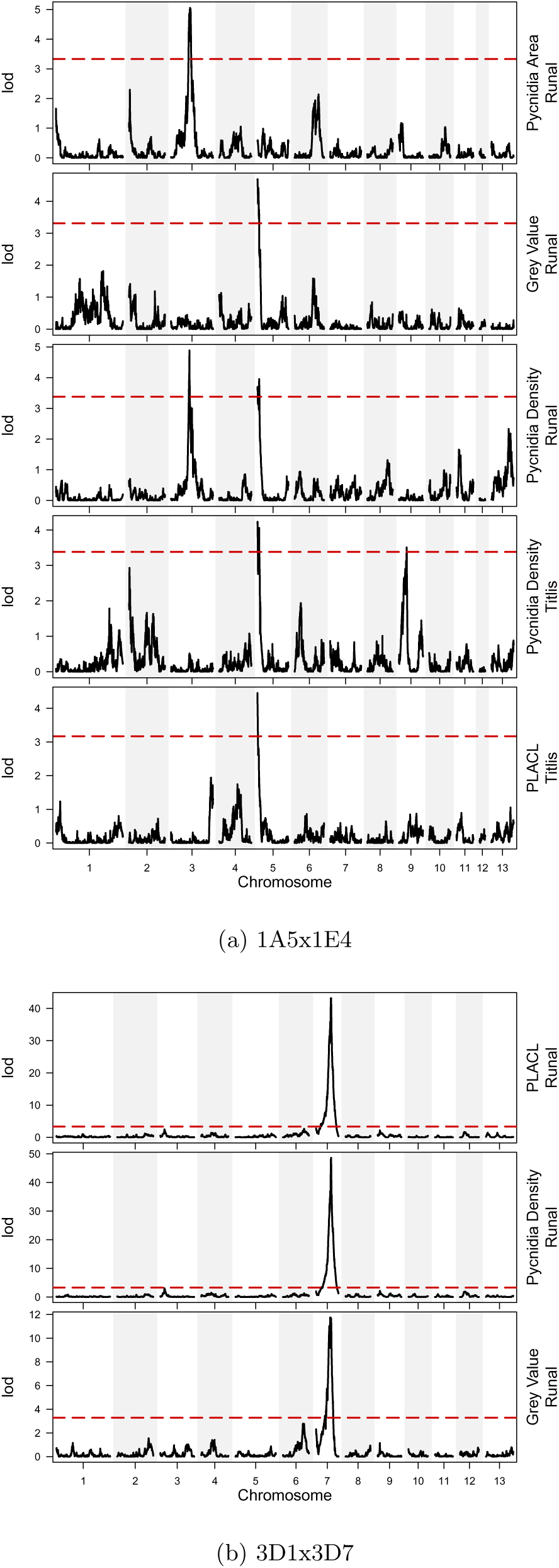
LOD (logarithm of the odds) plots from QTL mapping of virulence traits in the *Zymoseptoria tritici* mapping populations 1A5×1E4 (a) and 3D1x 3D7 (b) for the 13 core chromosomes (x axis). The dashed horizontal line represents the 0.05 significance threshold calculated with 1000 permutations.

The single, large effect QTL on chromosome 7 in cross 3D1×3D7 explained 54% of the phenotypic variance for PLACL, 57% of the variance for pycnidia density and 18% of the variance for pycnidia grey value. PLACL and pycnidia density shared the same confidence interval containing 35 genes. The wider confidence interval for pycnidia grey value contained 227 genes (Table 1A). The allele from the 3D7 parent was responsible for higher PLACL and pycnidia density and decreased pycnidia melanization.

**Table 1.1:** Significant QTLs for in planta virulence traits from *Zymoseptoria tritici* mapping populations 3D1x3D7 (A) and 1A5×1E4 (B) on the wheat cultivars Runal and Titlis.

| Cultivar | Phenotype | Chr | LOD | Variance Explained (%) | LOD Threshold | p value | Confidence Interval (Mb) | Number of Genes | Mean Phenotype (3D1 allele) | Mean Phenotype (3D7 allele) |
| --- | --- | --- | --- | --- | --- | --- | --- | --- | --- | --- |
| Runal | PLACL | 7 | 43.7 | 53.9 | 3.34 | <0.001 | 1.75-1.91 | 35 | 38.98 | 78.19 |
| Runal | Pycnidia per cm<sup>2</sup> | 7 | 48.2 | 57.3 | 3.3 | <0.001 | 1.75-1.91 | 35 | 11.26 | 34.23 |
| Runal | Melanization | 7 | 11.4 | 18.1 | 3.29 | <0.001 | 1.18-1.91 | 227 | 71.72 | 78.44 |

Table 1.1: Significant QTLs for in planta virulence traits from *Zymoseptoria tritici* mapping populations 3D1x3D7 (**A**) and 1A5x1E4 (**B**) on the wheat cultivars Runal and Titlis.
| Cultivar | Phenotype | Chr | LOD | Variance Explained (%) | LOD Threshold | p value | Confidence Interval (Mb) | Number of Genes | Mean Phenotype (1A5 allele) | Mean Phenotype (1E4 allele) |
| --- | --- | --- | --- | --- | --- | --- | --- | --- | --- | --- |
| Runal | Pycnidia per cm<sup>2</sup> | 3 | 4.9 | 8.6 | 3.38 | 0.003 | 1.27-1.87 | 209 | 24.79 | 32.13 |
| Runal | Pycnidia per cm<sup>2</sup> | 5 | 3.8 | 7 | 3.38 | 0.018 | 0.14-0.23 | 280 | 31.02 | 26.60 |
| Runal | Pycnidia Size (mm<sup>2</sup> x 10<sup>-3</sup>) | 3 | 5.1 | 8.7 | 3.33 | 0.001 | 1.27-1.98 | 143 | 4.773 | 4.998 |
| Runal | Melanization <sup>a</sup> | 5 | 4.7 | 8.1 | 3.31 | 0.001 | 0.14-0.15 | 89 | 88.06 | 84.74 |
| Titlis | Pycnidia per cm<sup>2</sup> | 5 | 4.15 | 7.2 | 3.38 | 0.006 | 1.36-2.27 | 280 | 28.03 | 22.62 |
| Titlis | Pycnidia per cm<sup>2</sup> | 9 | 3.51 | 6.1 | 3.38 | 0.045 | 3.20-9.91 | 204 | 22.37 | 28.01 |
| Titlis | PLACL | 5 | 4.56 | 8 | 3.17 | 0.015 | 1.36-2.27 | 280 | 95.49 | 90.40 |

<sup>a</sup>The grey scale runs from 0 (black) to 255 (white). Smaller values denote darker pycnidia.

In cross 1A5×1E4 the variance explained by each QTL was smaller and ranged from 6.1% for pycnidia density on Titlis to 8.7% for pycnidia size on Runal. The number of genes within each confidence interval was generally higher than in cross 3D1×3D7, ranging from 89 for grey value in Runal to 280 in the QTL on chromosome 5 for pycnidia density in Runal and PLACL in Titlis (Table 1B). The 1A5 allele was responsible for the higher phenotype values in the QTL on chromosome 5 whereas the 1E4 allele was responsible for the higher phenotype values in the chromosome 3 and 9 QTLs.

Additive effects were observed for the two QTLs found for pycnidia density in 1A5×1E4. On Runal, isolates with the 1A5 allele at the chromosome 5 QTL peak and the 1E4 allele at the chromosome 3 QTL peak had significantly higher pycnidia density than isolates with the reverse alleles (F=11.2, p<0.001) (Table 2A). On Titlis, isolates with the 1E4 allele at the chromosome 9 QTL peak and the 1A5 allele at the chromosome 5 QTL peak had significantly higher phenotypes than isolates with the opposite allele combinations (F=19.1, p<0.001) (Table 2B). On both cultivars, isolates with the same parental allele at both QTL peaks were not different from each other and were intermediate to the isolates with the alleles coming from different parents.

**Table 1.2:** Allele effects for pycnidia density in cross 1A5×1E4 for cultivars Runal (A) and Titlis (B). Chromosome alleles represent the parental allele of the marker at the QTL peak. Phenotypes are the mean pycnidia density of all isolates with the corresponding allele combination. Superscript letters denote significant differences. *n* represents the number of isolates with each allele combination.

| Cultivar | Chr 3 allele | Chr 5 allele | Mean Phenotype | *n* |
| --- | --- | --- | --- | --- |
| Runal | 1E4 | 1A5 | 39.28<sup>a</sup> | 43 |
| Runal | 1A5 | 1A5 | 26.10<sup>b</sup> | 32 |
| Runal | 1E4 | 1E4 | 26.96<sup>b</sup> | 44 |
| Runal | 1A5 | 1E4 | 23.09<sup>c</sup> | 46 |

Table 1.2: Allele effects for pycnidia density in cross 1A5x1E4 for cultivars Runal (A) and Titlis (B).
| Cultivar | Chr 5 allele | Chr 9 allele | Mean Phenotype | *n* |
| --- | --- | --- | --- | --- |
| Titlis | 1A5 | 1E4 | 33.38<sup>a</sup> | 37 |
| Titlis | 1A5 | 1A5 | 25.00<sup>b</sup> | 41 |
| Titlis | 1E4 | 1E4 | 25.15<sup>b</sup> | 54 |
| Titlis | 1E4 | 1A5 | 20.11<sup>c</sup> | 36 |

### Effect of Accessory Chromosomes on Virulence

Among the parent isolates used to make the crosses, all accessory chromosomes were present in 3D1 whereas 3D7 was missing chromosomes 14, 15, 18 and 21. In the 1A5×1E4 cross chromosome 17 was absent in 1E4. QTLs could only be mapped on chromosomes present in both parent isolates. No QTLs were found on the accessory chromosomes in either cross.

The presence or absence of accessory chromosomes was established for each of the progeny in cross 3D1×3D7. Chromosome 14 was absent in 67 (26%) progeny, chromosome 15 in 68 (26%) progeny, chromosome 18 in 77 (30%) progeny and chromosome 21 in 73 (28%) progeny. All accessory chromosomes were present at a significantly higher frequency than the expected 1:1 Mendelian inheritance ratio (chromosome 14: *χ*^2^=61.1, p<0.001, chromosome 15: *χ*^2^= 59.1, p<0.001, chromosome 18: *χ*^2^ = 43.1, p <0.001, chromosome 21: *χ*^2^ = 49.9, p<0.001). This is consistent with previous work that reported skewed inheritance of accessory chromosomes in a subset of 48 isolates from the same population (Croll et al. 2013).

We studied the effect of accessory chromosome presence-absence on virulence. On Runal, isolates with chromosome 21 had significantly larger pycnidia (F=6.2, p=0.014) than isolates without chromosome 21, but the effect size was small (*η*^2^=0.023). On Titlis, isolates with chromosome 18 had significantly larger pycnidia (F=5.0, p=0.027) and darker pycnidia (F=5.7, p=0.018) than isolates without chromosome 18. The effect sizes for pycnidia size (*η*^2^=0.021) and pycnidia melanisation (*η*^2^=0.022) were small. A significant interaction between chromosomes 15 and 18 was also observed for pycnidia size (F=7.0, p= 0.009), with isolates having both accessory chromosomes showing larger pycnidia, but the effect size of the interaction was small (*η*^2^=0.028).

### Candidate genes for pathogen virulence within QTLs

In total, 918 candidate genes were identified in the 95% confidence intervals of all QTLs, including 227 from cross 3D1×3D7 and 691 from cross 1A5×1E4. No candidate genes were shared between the two crosses. In the literature to date, 19 genes have been implicated in virulence in *Z. tritici* and functionally characterized (Table S1). None of the genes previously shown to play a role in virulence were found within the QTLs.

### In-depth analysis of the chromosome 7 QTL

The QTL for PLACL and pycnidia density in cross 3D1×3D7 on Runal had a high LOD score, a low number of candidate genes within the 95% confidence interval, and explained approximately half of the observed phenotypic variance. Existing gene models were manually checked for accuracy, sequence polymorphisms between the two parent isolates were analyzed and gene expression during the infection cycle was investigated. Population genomic approaches were used to investigate the genomic features of the region in a natural population.

### Re-annotation of the genes in the chromosome 7 QTL

Within the 95% confidence interval, 25 of the 35 gene models from the reference annotation were not convincingly supported by RNAseq reads. The annotations of 20 genes were replaced with those of Grandaubert et al. (2015). Four genes not present in any previous annotation were discovered and supported by our RNAseq data. The annotations of four additional genes were altered slightly from those of Grandaubert et al. (2015) or the JGI. After re-annotation, 38 genes were found in the 95% confidence interval (Table 3, Table S2). Among the newly annotated genes, GO terms could be assigned to 17 genes while the remaining 21 genes had no known function. Six genes within the QTL encode proteins with a predicted signal peptide but lack a trans-membrane domain and are therefore predicted to be secreted. Among these six genes, two meet the criteria for being small secreted effector-like proteins (<300 amino acids, secreted and cysteine rich).

**Table 1.3:**
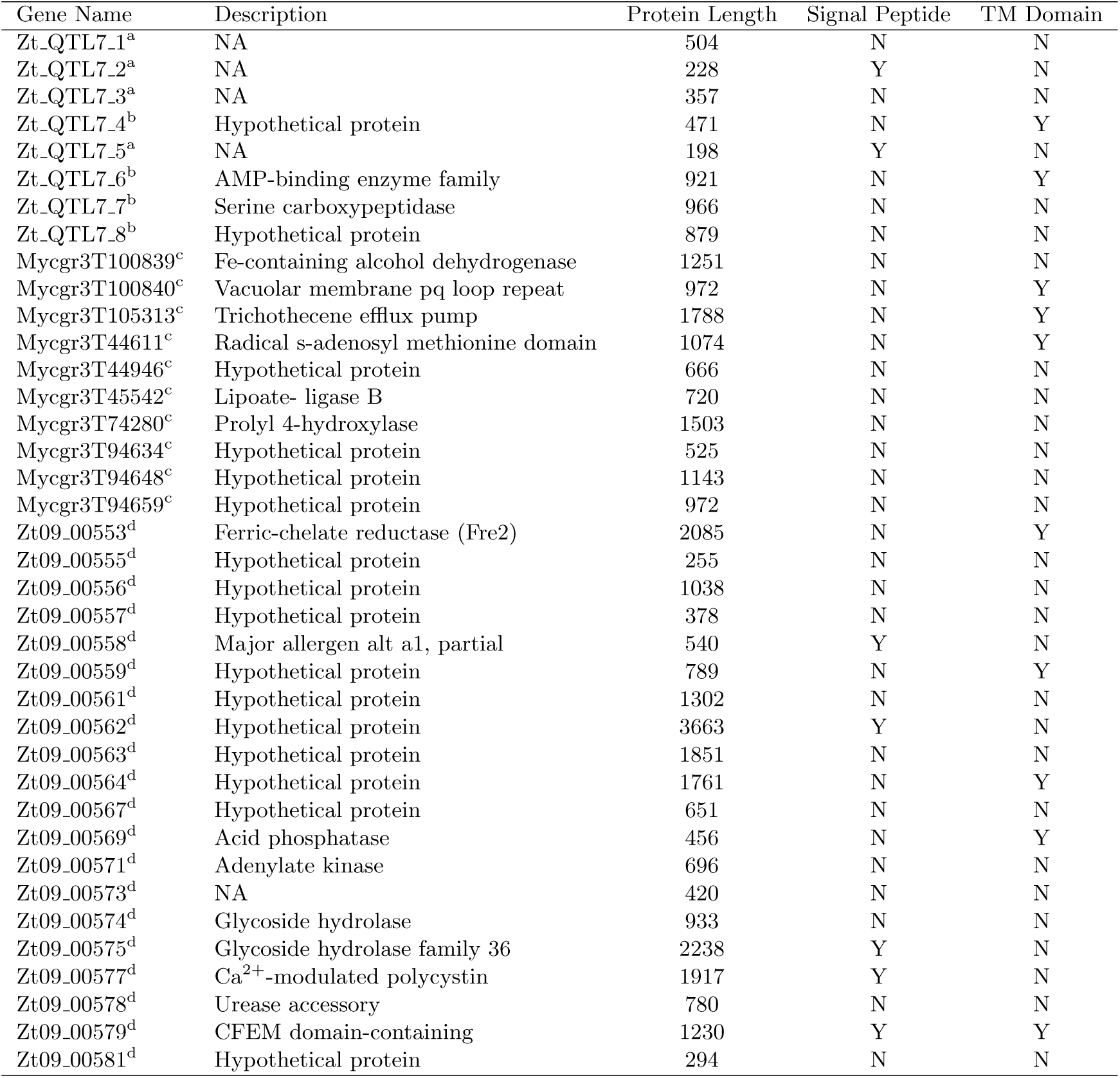
Re-annotated candidate genes located in a QTL for pycnidia density and PLACL on chromosome 7 in *Zymoseptoria tritici*. Superscripts denote the source of the gene annotation: ^a^new genes not present in previous annotations, ^b^ annotations modified from existing annotations, ^c^ annotation from the JGI reference annotation, ^d^ annotation from Grandaubert et al. 2015. Gene descriptions are GO terms from blast2go. Protein length is the number of amino acids in the mature protein. Presence (Y) and absence (N) of signal peptides and trans membrane (TM) domains are indicated.

### Identification of high priority candidate genes within the chromosome 7 QTL

Sequence variation within gene coding regions of the chromosome 7 QTL was investigated in the parental genomes. 82 sequence variants were identified, comprising 51 synonymous and 31 non-synonymous mutations as well as 4 insertions/deletions (indels). 17 genes were found to have at least one mutation affecting amino acid sequence. 19 genes showed differential expression in planta between the two parental isolates for at least one time point during the infection process. Several high-priority candidate genes were identified based on putative function, sequence variation and expression levels during infection. Candidate gene Zt_QTL7_5 is not present in any existing annotation and BLAST searches showed no similarities to other proteins. With a length of 65 aa, six cysteine residues and a signal peptide domain, it fulfills the criteria of a typical small secreted effector protein. Its peak expression is between 12 and 14 dpi, during the transition from symptomless growth to the onset of chlorosis (Figure 4). It is the second most expressed gene in the QTL confidence interval, but shows significantly lower expression in 3D7 (the more virulent parent) at 14 dpi. Candidate gene 00558 is also a small secreted protein containing an *Alternaria alternata* allergen domain. This domain is unique to the Dothideomycete and Sordariomycete classes of fungi (Chruszcz et al. 2012) and can induce major allergic reactions in the human respiratory systems (Bush et al. 2001). This gene shows peak expression during the symptomless phase and lower expression in 3D7 (Figure 4). Candidate gene 105313 is a fungal-specific transporter from the Major Facilitator Superfamily (MFS). It contains a trichothecene efflux domain that has been shown to play a role in toxin secretion in Fusarium (Alexander et al. 1999). The parental alleles differed by two non-synonymous SNPs. Candidate gene 00579 is a fungal-specific membrane protein containing 7 trans-membrane domains, a CFEM domain and a signal peptide. Expression peaks at 7 dpi (Figure 4).

**Figure 1.4:**
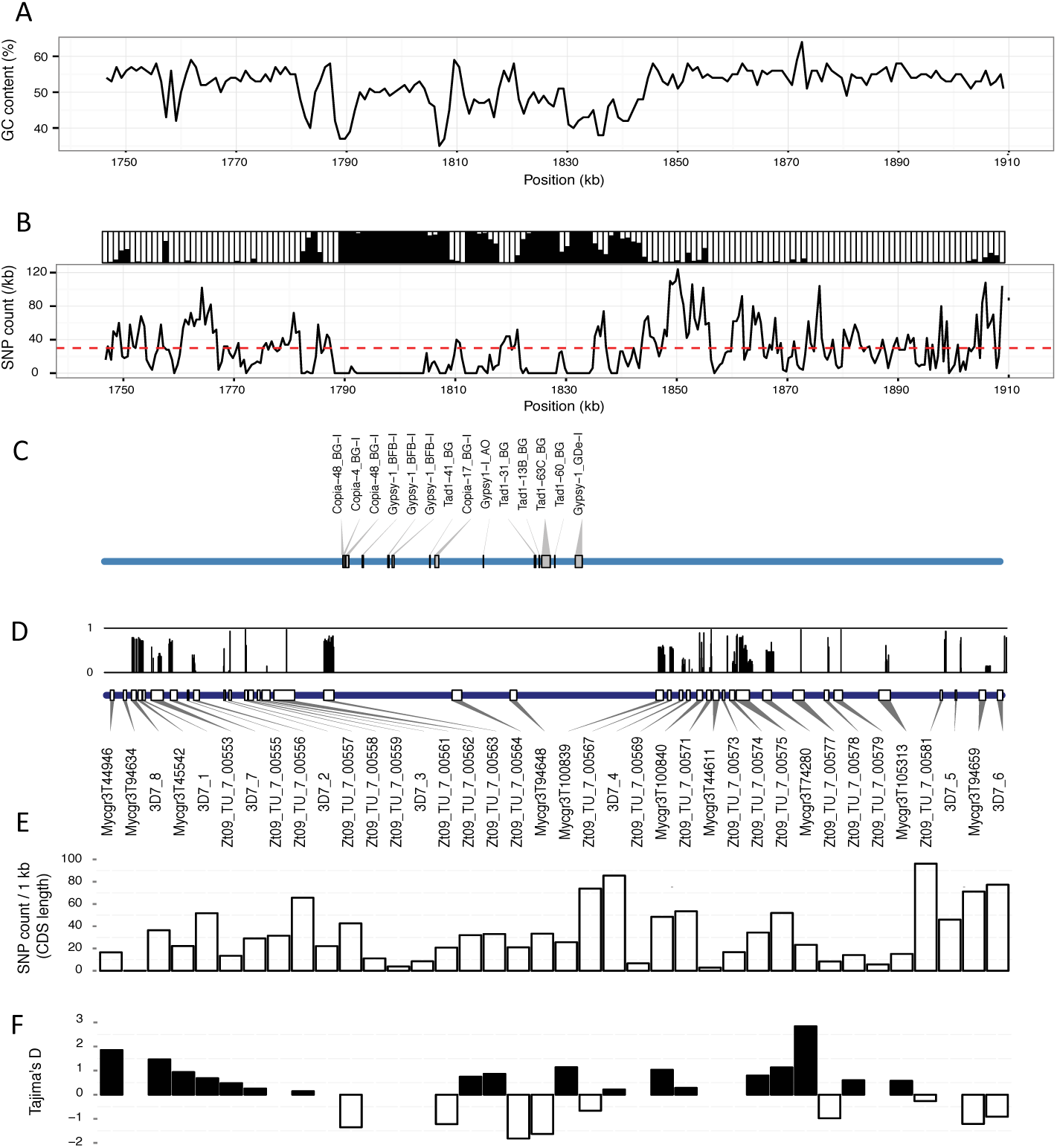
Population genomic characterisation of the QTL located on chromosome 7 (region 1746 kb to 1909 kb) in the reference isolate IPO323 and a population of 28 *Zymoseptoria tritici* field isolates from Switzerland. (A). GC content in the IPO323 reference genome. (B). SNPs density (bottom) and genotyping rate (top). Black rectangles represent the frequency of missing SNP genotypes. The horizontal line represents the mean SNP density across chromosome 7 in the 28 field isolates. (C). Transposable element locations in the IPO323 reference genome. (D). Bottom: Location of coding sequences in the IPO323 reference genome. Top: Frequency of the allele carried by 3D7 in the 28 field isolates for the SNPs shared by the 3D1 and 3D7 parents in the coding sequences of the predicted genes. (E). SNP density per 1 kb in the coding sequences of the predicted genes. (F). Tajima’s D statistic per gene coding sequence. Tajima’s D values for genes with less than 10 SNPs are not shown.

### Population genomics of genes in the chromosome 7 QTL

As QTL mapping is highly cross specific, genetic variation in the chromosome 7 QTL was investigated in a natural population of 28 re-sequenced *Z. tritici* isolates from Switzerland. The 163 kb confidence interval contained an alternation of gene-rich and gene-poor regions. A large gene-poor region (found between ca. 1.79 - 1.84 Mb in the reference genome assembly) contained only two genes (Figure 3E). This gene-poor region contained all transposable elements (TEs) found within the confidence interval (Figure 3D). Three classes of TEs were found in this region, including Copia, Gypsy and Tad1 elements. The gene-rich regions contained a higher density of SNPs than the average on the chromosome, reaching up to 120 SNPs per kb in some regions (Figure 3C). The genotyping rate was high in the gene-rich region. The gene-poor and TE-rich regions contained few SNPs and had a low genotyping rate (Figure 3C) and generally had a lower GC content than the gene-rich regions (Figure 3B).

**Figure 1.3:**
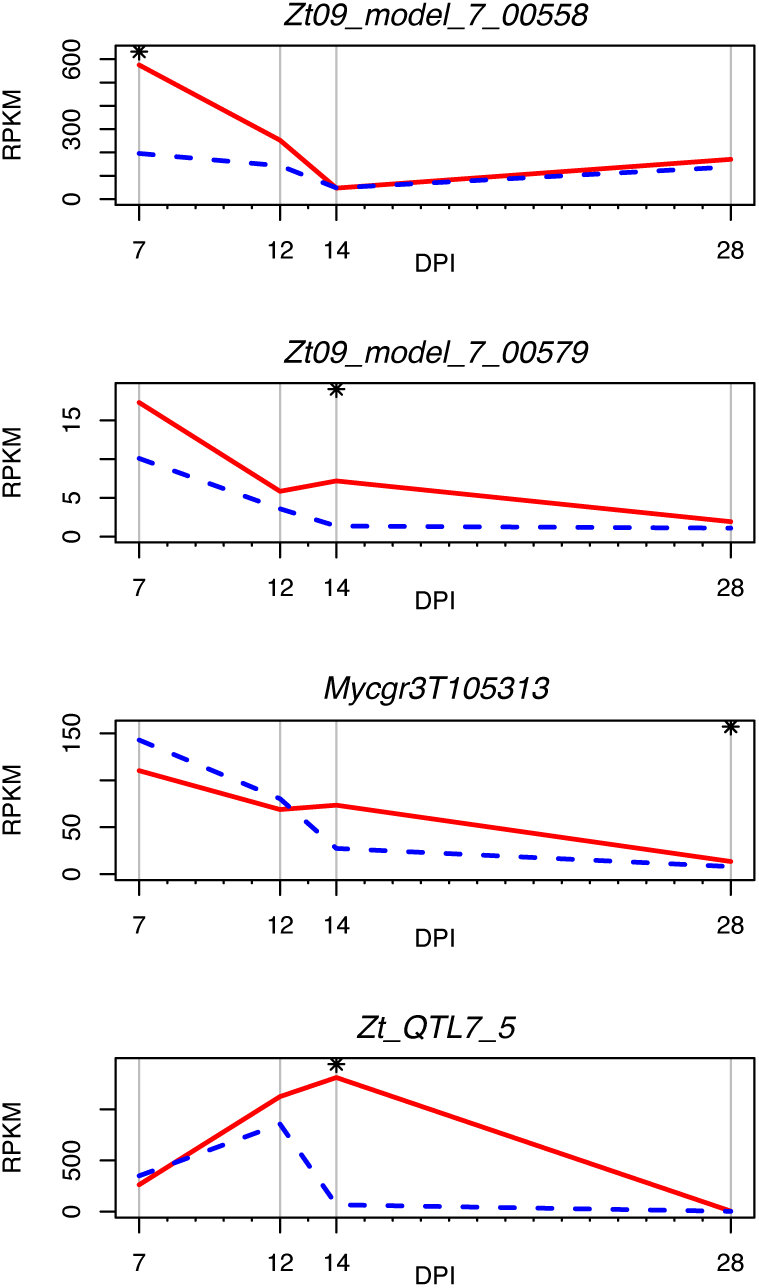
Gene expression profiles of *Zymoseptoria tritici* candidate genes at 7, 12, 14 and 28 days post infection (DPI). Solid red lines represent parent isolate 3D1, dashed blue lines represent parent isolate 3D7.

Genes within the confidence interval were highly variable for the number of SNPs within the natural population (Figure 3F). Seven genes contained fewer than 10 SNPs per kb whereas 9 genes contained more then 50 SNPs per kb. On average, the genes within the confidence interval contained 33 SNPs per kb. The Tajima’s D statistic for 27 genes having 10 or more SNPs showed an alternation of clusters of genes with positive D values and genes with negative D values (Figure 3G). Genes with negative D values are more likely to be under purifying selection while genes with positive D values are likely to be under balancing selection. This suggests variability in the selection processes acting on these genes, including a possibility of hitchhiking effects.

## Discussion

Here we report the most comprehensive QTL mapping analysis of fungal virulence to date, utilizing different mapping populations and different host cultivars as well as measuring multiple traits associated with virulence. We were able to genetically dissect pathogen virulence into separate components, showing that different components of virulence were under separate genetic control. QTLs were identified for all traits, including QTLs that were specific to mapping populations, cultivars and traits as well as QTLs that were shared among traits within the same mapping population. We identified genome regions associated with single large effects as well as regions of smaller additive effects.

### The genetic architecture of virulence in *Z. tritici* is complex

Our results point to a complex genetic architecture involving multiple factors that combine to produce quantitative virulence. Strong evidence was found for transgressive segregation, with many progeny showing more extreme phenotypes than the parent isolates in both crosses. QTLs were specific to each cross and in some cases specific to a phenotype or cultivar. In all but one case, multiple traits mapped to the same QTL, an observation also made for other traits in *Z. tritici* (Lendenmann et al. 2014; Lendenmann et al. 2015; Lendenmann et al. 2016).

Both quantitative (Zhan et al. 2005) and large-effect interactions consistent with the gene-for-gene (GFG) hypothesis have been reported in *Z. tritici* (Brading et al. 2002; Kema et al. 2000). GFG interactions are frequently reported as conferring compatible/incompatible interactions (Flor 1955). The large effect size of the 3D1×3D7 chromosome 7 QTL on one of the cultivars shows similarity to a GFG interaction between a major resistance gene in Runal and a corresponding avirulence effector gene in the 3D1 strain of the pathogen. In our case the interaction explains ^˜^50% of the overall variance for the virulence traits which also show a continuous distribution in the mapping population. These observations suggest that numerous additional genes of small effect may also be involved, in addition to the genotype-by-environment interactions typically observed in quantitative phenotypes.

In contrast to the single large effect QTL in 3D1×3D7, numerous QTLs distributed across several chromosomes were found for the different phenotypes in cross 1A5×1E4. These QTLs had relatively small effects on phenotype and contained a high number of candidate genes. This suggests that the genetic architecture of virulence in 1A5×1E4 is more complex than that of 3D1×3D7 and demonstrates the contribution of both large and small effects on virulence.

### Additive effects of QTL loci on virulence

The multiple factor hypothesis (Morgan et al. 1915) based on additive effects of many genes is the key principle underlying quantitative traits. Recombination generates progeny with different combinations of genes than the parents and can contribute to novel virulence phenotypes (Joseph et al. 2011; Kanvil et al. 2015). Strong additive effects were observed in cross 1A5×1E4 for the QTL for pycnidia density. For both cultivars, alternative alleles from the two parents combined to produce a higher phenotype. Isolates with the same parental allele at both QTLs showed lower virulence than isolates with the different parental alleles at the two QTLs. Transgressive segregation was observed for all phenotypes in both crosses. These findings illustrate the power of recombination to generate progeny phenotypes that are significantly different from their parents, an observation shared with other plant pathogenic fungi (Sommerhalder et al. 2010; Stefansson et al. 2014) and show how a range of quantitative virulence phenotypes can be generated through sexual reproduction in natural field populations.

### The contribution of accessory chromosomes to virulence

By using the novel approach of associating virulence phenotypes with the presence or absence of specific accessory chromosomes, we discovered a significant correlation for some phenotypes, with higher virulence observed in offspring carrying certain accessory chromosomes. Accessory chromosomes were shown to play an important role in virulence in other fungal pathogens (Akagi et al. 2009; Ma et al. 2010; Miao et al. 1991) but until now their role in *Z. tritici* was elusive (Stukenbrock et al. 2010). The increase in virulence was small, around 2-3%, but significant. If this correlation also exists under natural field conditions, this increase in virulence and the associated increase in reproduction would represent a significant fitness advantage that could explain why accessory chromosomes are retained in natural field populations.

### Cultivar specificity

Cultivar specificity is well documented in *Z. tritici* (Ahmed et al. 1995; Cowger et al. 2002; Zhan et al. 2002) and is commonly reported as qualitative compatible/incompatible interactions (e.g. (Brading et al. 2002)) or as generally quantitative with isolates being more virulent or less virulent on a particular host (e.g. (Zhan et al. 2005)). In cross 3D1×3D7, all phenotypes were higher on the more susceptible cultivar as expected. However, in cross 1A5×1E4, PLACL was higher in Runal whereas pycnidia size, density and melanization were higher in Titlis, the more resistant cultivar. Additionally, in 1A5×1E4 the same chromosome 5 QTL was found for pycnidia density in both cultivars. However, a second QTL was found on chromosome 3 in Runal and on chromosome 9 in Titlis, suggesting that these QTL loci play a role in quantitative specialization to these two cultivars. This is in contrast to the findings of Mirzadi Gohari et al. 2015 who found a QTL at a similar position on chromosome 5 that was implicated in host specificity in a different set of cultivars. The results from the accessory chromosome presence/absence analysis also point towards cultivar specificity, with different accessory chromosomes affecting the same phenotype on different cultivars.

### Contribution of pycnidia number and size to epidemic potential

The basic reproductive number (R_0_) is the theoretical number of infections arising from a single infection event and can be useful in predicting the potential development of plant disease epidemics (van den Bosch et al, 2008). Pycnidia represent the asexual reproductive output of a *Z. tritici* strain, with larger pycnidia correlated with a greater number of conidia in earlier work (Gough, 1978). It was previously postulated that the number and size of pycnidia could be important measures of virulence in *Z. tritici* (Stewart et al. 2014; Suffert et al. 2013). Our results show a significant correlation indicating that larger pycnidia bear both larger and more numerous spores.

The number and size of pycnidia influence the R_0_ and therefore provide a prediction of the epidemic potential associated with a pathogen strain under field conditions. In contrast, measuring the leaf area covered by lesions (PLACL) gives an indication of the amount of damage caused by a pathogen strain on the host plant. The ability to directly measure both the epidemic potential of an isolate and the amount of damage it causes to its host represents a powerful improvement in the overall measurement of pathogen virulence that can also be used to improve measurements of resistance in the host (Stewart et al. 2016). Furthermore, the identification of a QTL for pycnidia size shows that this important trait can be under separate genetic control. A separate experiment oriented around measuring differences in host resistance and conducted using naturally infected plants under field conditions showed that host genotype could affect pycnidia formation independently of damage caused by leaf lesions (Stewart et al. 2016). Theoretically, a pathogen strain which causes high levels of leaf necrosis but has a low R_0_ (i.e. producing fewer pycnidia per infected leaf) may appear more damaging at a small spatial scale but an isolate which has a higher R_0_ (i.e. producing more pycnidia per infected leaf) has a greater potential to cause damage over larger spatial scales. From an epidemiology perspective, directly measuring pathogen reproduction on a given host could inform disease management decisions oriented around inhibiting pathogen reproduction (e.g. through seeking resistance genes that lower pathogen reproduction in addition to resistance genes that lower host damage), as well as providing a novel way of selecting for quantitative host resistance. The results from cross 1A5×1E4 illustrate the potential usefulness of this approach with different levels of pycnidia and PLACL found on the two different cultivars.

### Candidate genes affecting pathogen virulence

By combining several disciplines, we were able to narrow the candidate gene list for the chromosome 7 QTL affecting PLACL, pycnidia density and melanization, but it remains possible that different genes located within this QTL interact to affect all three traits. It also is possible that one gene in this QTL is primarily responsible for only one phenotype, which influences the other two without any genetic or regulatory connection, a situation referred to as spurious, vertical, relational or reactive pleiotropy (Paaby et al. 2013). For example, it is plausible that a gene encoding higher PLACL would lead to greater pycnidia density that could in turn affect pycnidia melanization. The results from cross 1A5×1E4 offer support for the latter hypothesis, with multiple traits mapping to the same QTL on chromosome 5, though independent QTLs were found on different chromosomes for the same traits. In previous work, genes up-regulated in the biotrophic phase of the infection cycle showed reduced pycnidia production in knockout mutants (Poppe et al. 2015), providing evidence that pycnidia development is somewhat dependent on prior processes.

Among the candidate genes within the chromosome 7 QTL, the one we consider most likely to explain the observed virulence is Zt_QTL7_5, a previously un-described small secreted protein (SSP) that is highly expressed during the switch to necrotrophic growth. SSPs are common virulence factors in fungi that facilitate infection or elicit a response in the host (Lo Presti et al. 2015). SSPs often conform to the GFG paradigm by interacting with a major R gene in the host. A GFG interaction would explain the large effect of the chromosome 7 QTL on only one host. Interestingly, no significant sequence differences exist for this gene between the parent isolates, but the expression levels are significantly lower in 3D7, the more virulent parent. This is consistent with the theory of effector triggered immunity (ETI), where recognition of a pathogen effector triggers a defense response in the host. Another SSP, candidate gene 00558, contains a conserved domain known to elicit allergic response in humans (Bush et al. 2001). It has additionally been implicated in virulence in *Alternaria brasicicola* on Arabidopsis (Cramer et al. 2004) and inhibits plant antimicrobial proteins (Gómez-Casado et al. 2014). Given the existing knowledge of this protein, it appears that it could play a role in *Z. tritici* virulence. An MFS transporter implicated in toxin secretion was also located in the QTL. MFS transporters contribute to virulence by secretion of fungal toxins and secondary metabolites as well as efflux of plant-derived antimicrobial compounds (Coleman et al. 2009). Candidate gene 00579 contains a CFEM domain that is unique to fungi and found at significantly higher frequency in pathogenic than non-pathogenic fungi (Zhang et al. 2015). Putative functions of the CFEM domain include cell surface receptors, signal transduction or adhesion of molecules in plant-pathogen interactions (Kulkarni et al. 2003). None of the genes previously described as having a role in virulence in *Z. tritici* were identified in this study, highlighting the usefulness of forward genetic approaches such as QTL mapping to identify novel virulence genes.

### Population genomics of the chromosome 7 QTL

Analysis of the large effect QTL on chromosome 7 in a Swiss field population revealed several interesting genomic features. The region was highly variable, with genes exhibiting variable numbers of SNPs as well as evidence for both purifying and balancing selection. Genes involved in virulence are often under positive selection in plant pathogens (Lo Presti et al. 2015; Stukenbrock et al. 2009), including in *Z. tritici* (Poppe et al. 2015). An island of transposable elements (TEs) was found within a gene-poor region of the QTL. The *Z. tritici* genome contains 16.7% repetitive elements (Dhillon et al. 2014). Effector genes show a tendency to cluster in gene-poor regions of repetitive DNA that are TE-rich in numerous filamentous fungal plant pathogens (Dong et al. 2015). TEs can influence the expression of effector genes by insertion into promoter regions (Ali et al. 2014) or through epigenetic gene silencing (Shaaban et al. 2010). We hypothesize that these processes may explain the differences in expression observed in *Zt_QTL7_5*.

### Conclusions

This study highlights the complex nature of virulence in the wheat-*Z. tritici* pathosystem, illustrating that many factors contribute to quantitative phenotypes. We showed that virulence is comprised of different traits, some affecting host damage and others affecting pathogen reproduction, that can be under independent genetic control. In light of this, researchers and breeders should reconsider the best way to measure virulence in this pathosystem. We propose that more attention should be focused on resistance that reduces pathogen reproduction in order to decrease R_0_ during epidemics.

## Experimental Procedures

Two *Z. tritici* mapping populations (described in Lendenmann et al. 2014) were phenotyped in a greenhouse-based seedling assay as described in Stewart et al. 2014. The Swiss winter wheat cultivars ‘Runal’ and ‘Titlis’ (DSP Ltd, Delly, Switzerland) were inoculated using each offspring from both crosses along with the parental isolates. Two plants of each cultivar were inoculated with each *Z. tritici* isolate to give two technical replicates. Each technical replicate was placed in a separate greenhouse compartment. This process was repeated three times over three consecutive weeks to generate three biological replicates, resulting in six replicates total for each isolate-cultivar combination.

At 23 days post inoculation (dpi) the second leaf from each plant was excised, photographed and phenotyped using automated image analysis as described previously (Stewart et al. 2014). The method was modified slightly to include a measure of pycnidia melanization. RGB images were converted to 8-bit greyscale and the mean grey value for the pixels making up each pycnidium was calculated. The grey scale runs from 0 (black) to 255 (white). Grey values can be used as a proxy for the degree of melanisation (Lendenmann et al. 2015). The phenotypes percent leaf area covered by lesions (PLACL), pycnidia density (pycnidia per cm^2^ leaf area), pycnidia size (in mm^2^) and pycnidia melanization were used as phenotypes for QTL mapping (see below).

For the 3D1×3D7 progeny, the mean pycnidia size was calculated for each isolate on Runal. The 10 isolates showing the largest mean pycnidia sizes and 10 showing the smallest mean pycnidia sizes were selected for further analysis. The leaves from the six replicates for each isolate were retrieved from storage. Each leaf was cut into ^˜^30 mm long sections. All sections from each leaf were placed into a 1.2 ml collection microtube (QIAGEN) along with a 5 mm x 25 mm strip of filter paper. 100 *μ*l sterile water was added to each tube to moisten the filter paper. Tubes were capped and placed at 25°C for 24 h to provide the humid environment needed to exude the cirrhi containing spores from pycnidia. 800 *μ*l of water containing 0.001% TWEEN 20 was added to each tube. Tubes were vortexed for 20 seconds to suspend the released spores. 100 *μ*l of the spore solution was placed into a black glass-bottomed 96 well plate (Greiner bio one, *μ*Clear). Spores were imaged with an Olympus IX 81 inverted microscope coupled with a Hamamatsu ORCA-ER camera using transmission illumination. Four images were made from each well at 20x magnification with a 10% overlap between images. Images from each well were stitched together using ImageJ (Rasband, 1997-2015). Each spore was counted and its length was measured using the line selection tool and the measure command in ImageJ (Figure S1). The mean spore length was used as a proxy for spore size. The total number of spores per leaf was calculated (Figure S1) and divided by the number of pycnidia per leaf to derive the mean number of spores per pycnidium.

All analyses were performed in base R (R Core Team 2012) unless specified otherwise. To normalize environmental variation between technical replicates and biological replicates and account for genotype-by-environment (GxE) interactions, phenotypes were mean centered (Schielzeth 2010). For each phenotype, the mean value for each greenhouse chamber and time point was calculated. This value was subtracted from all individual values from the corresponding greenhouse chamber and time point, resulting in a mean value of ^˜^0 for each greenhouse chamber time point. The mean of these transformed values was calculated for each progeny and used for subsequent analyses. PLACL and pycnidia density values were log transformed prior to mean centering. Mean centered values were used for analysis but untransformed values are reported in the text and figures for clarity.

Phenotype differences between wheat cultivars were calculated using ANOVA. Effects on phenotypes of accessory chromosome presence or absence were calculated using ANOVA with the presence or absence of each chromosome as factors. Non-significant factors were removed from the models. Effect sizes (*η*^2^) were calculated using the lsr package (Navaro 2015). Fisher’s LSD test was used to calculate the differences between isolates with different parental alleles at QTL peaks using the Agricolae package (deMendiburu 2014). Correlations between pycnidia size and spore size and between pycnidia size and number of spores per pycnidia were made with Pearson’s correlation coefficient.

Progeny from both crosses were genotyped using restriction site associated DNA sequencing (RADseq) based on a method adapted from Etter et al. 2011 as described previously (Lendenmann et al. 2014). The genetic map of Lendenmann et al. 2014 was used to perform QTL mapping. QTL mapping was performed using the r/QTL package (Arends et al. 2010) following the methods described in Lendenmann et al. 2014.

To establish the presence or absence of accessory chromosomes in 3D1×3D7, sequence reads from the RADseq genotyping were aligned to the IPO323 reference genome (Goodwin et al. 2011). For each isolate, the mean read depth was calculated for each chromosome and divided by the mean sequence depth over all chromosomes. Thus, chromosomes with read depth similar to the genome average have a value of ^˜^1 and were deemed present whereas chromosomes with low or no read depth have a value of ^˜^0 and were deemed absent. Deviations from Mendelian inheritance among the accessory chromosomes exhibiting presence/absence polymorphisms in the parents were tested using a Chi square (*χ*^2^) test.

Parent genome sequences (Croll et al. 2013) were aligned to the IPO323 reference genome. Sequence variants were identified using the unified genotyper within GATK (McKenna et al. 2010) and effects of variants found within genes were predicted using snpEFF (Cingolani et al. 2012b) and snpSIFT (Cingolani et al. 2012a) following the methods outlined in Lendenmann et al (2014).

The genes within the 95% confidence interval containing a QTL with large effect on chromosome 7 were manually re-annotated using existing annotations from the IPO323 reference annotation (Goodwin et al. 2011) and Grandaubert et al. 2015 along with RNAseq data from IPO323 (Rudd et al. 2015) and from the 3D7 parent isolate (Palma-Guerrero et al. 2016). Genes were annotated using blast2go (Conesa et al. 2005). Signal peptides were identified using signalP 4.1 (Petersen et al. 2011) and trans-membrane domains were identified with TMHMM 2.0 (Krogh et al. 2001). The re-annotated genes were used for all subsequent analyses of this region.

An *in planta* infection assay was performed with the parent isolates 3D1 and 3D7 (Javier Palma-Guerrero, unpublished). RNA was sequenced from leaf samples collected at 7, 12, 14 and 28 days post inoculation (dpi) as described earlier (Palma-Guerrero et al. 2016).

The population genomics of the genes in the 95% confidence interval of the chromosome 7 QTL was characterized using 28 re-sequenced *Z. tritici* isolates sampled from a single field in Switzerland. Sequences were aligned to the reference genome using Bowtie 2 version 2.2.3 (Langmead et al. 2012). SNPs among the isolates were identified using GATK tools (DePristo et al. 2011). GC content was calculated in the reference genome using 1000 bp overlapping windows, with 100 bp overlaps. The locations of transposable elements within the reference genome were identified using RepeatMasker (Smit et al. 2015). The repeat library used for annotation was version 20150807 downloaded from Repbase in September 2015 (Jurka et al. 2005). The SNP genotyping rate in the 28 re-sequenced isolates was calculated in windows of 1000 bp. The SNP density was calculated in windows of 500 bp for SNPs that were genotyped in >50% of the re-sequenced isolates. The SNP density per coding sequence was calculated by dividing the number of SNPs with a genotyping rate ¿ 90% by the total coding sequence length. Tajima’s D statistic (Tajima 1989) was calculated per gene using Popgenome in R (Pfeifer et al. 2014). Only SNPs with a genotyping rate > 90% were included. Tajima’s D values for genes with less than 10 SNPs were not calculated.

## Acknowledgements

The research was supported by a grant from the Swiss National Science Foundation (31003A 134755). RADseq libraries were prepared at the Genetic Diversity Centre Zürich (GDC) and sequenced at the Quantitative Genomics Facility at the Department of Biosystems Science and Engineering (D-BSSE) at the scientific central facilities of ETH Zürich. Microscopy was carried out at the Scientific Centre for Optical and Electron Microscopy (ScopeM) of the ETH Zürich. Technical assistance was provided by Renato Guidon, Thomas Kreienbühl and Leandra Knecht. Cross 3D1×3D7 was made by M. Lendenmann. Cross 1A5×1E4 was made by Marcello Zala.

